# Transcription Factor Activation Profiles (TFAP) identify compounds promoting differentiation of Acute Myeloid Leukemia cell lines

**DOI:** 10.1101/2021.06.22.449489

**Authors:** Federica Riccio, Elisa Micarelli, Riccardo Secci, Giulio Giuliani, Simone Vumbaca, Giorgia Massacci, Luisa Castagnoli, Claudia Fuoco, Gianni Cesareni

## Abstract

Repurposing of drugs for new therapeutic use has received considerable attention for its potential to reduce time and cost of drug development. Here we present a new strategy to identify chemicals that are likely to induce differentiation of leukemic cells. As Acute Myeloid Leukemia (AML) is the result of a block in myeloid differentiation, finding new drugs that are capable of inducing blast terminal maturation is considered a valuable strategy. We used data from the Connectivity Map (CMap) to identify drugs that could be repositioned for their potential to activate transcription factors that mediate myeloid differentiation. Compounds promoting the activation of transcription factors that play a positive role in myeloid differentiation were considered candidate pro-differentiation drugs. This approach yielded a list of chemicals ranked according to the potential to activate transcription factors that induce differentiation of leukemic progenitor cells. Drugs that are already used in differentiation therapy, such as for instance all-trans retinoic acid (ATRA) are in the top positions of this ranked list. To validate our strategy, we tested the *in vitro* differentiation potential of 22 candidate compounds using the HL-60 human cell line as a myeloid differentiation model. Ten out of 22 compounds, ranking high in the inferred list, were confirmed to induce significant differentiation of HL-60. Some of these compounds are known to trigger the DNA damage response, thus identifying this process as a target to modulate myeloid differentiation. These results underscore the potential of our approach to accelerate the drug discovery process. The method that we have developed is highly versatile and it can be adapted to different drug repurposing projects.

## 1. Introduction

Pharmaceutical companies experience considerable hurdles in the development and marketing of new drugs [1]. This is because of the escalating cost and time required for preclinical research, clinical trials and regulatory requirements [2]. Thus, the process of drug development is today much slower than could have been anticipated from the rapid increase in our understanding of the molecular bases of human diseases. As a consequence, the identification of new therapeutic uses for drugs that are approved for different medical indications - process known as drug repositioning or repurposing - has attracted considerable interest as it allows acceleration of many of the required drug development steps [3]. Approximately 40 drugs are approved each year by the Food and Drug Administration (FDA) for different therapeutic uses and in 2019 over 20,000 chemicals or biologicals are validated for use in humans. This represents an impressive toolbox that can be used to modulate human physiology with minor undesired side effects. In principle a complete understanding of a pathology and a detailed annotation of each drug effects and side-effects should allow the design of new rational treatments to revert a disease phenotype. In practice such a direct approach is often not possible and the most successful cases of drug repurposing to date are the result of serendipitous observations and not of rational design or systematic approaches. Nevertheless, systematic approaches are still useful, as they can be applied on a large scale, and considerable effort is put in this direction. The validation of drugs that are candidates for repurposing requires labor-intensive preclinical procedures involving cell assays and testing on animal models before evaluation of efficacy in phase II clinical trials. Thus, the necessity to develop hypothesis generation approaches to identify drugs that are likely to be efficacious. This would limit the number of compounds that are worth considering for carrying over to more time-consuming steps. Different computational and experimental approaches have been developed for assisting in drug repurposing [3]. Computational approaches involve the integration of different data types to help formulate new hypotheses [4]. Gene expression profiles obtained by RNA-seq or proteomics approaches, genotype profiles, chemical structures and electronic health records have been used for this purpose [5].

The perturbation of the gene expression profile after drug treatment is a close proxy of the phenotypic changes caused by the drug on a cell or on a tissue. Since both drugs and mutations cause a perturbation of gene expression profiles, transcriptomics or proteomics data can be used to define and compare drug and disease signatures. Drugs causing similar gene expression changes (similar signatures) are likely to have similar phenotypic effect while drugs that have a signature that correlates negatively with the signature of a disease are candidates for “curing” the disease phenotype. This approach has been used with some success to group drugs according to signature or to identify candidates for reverting a disease phenotype [6]–[11]. Such a strategy has received a considerable boost from the Connectivity Map (CMap) project which yielded expression profiles and signatures for a large and growing number of cell states caused by chemical perturbations or mutations in different cell lines [12]. However, large scale generation of gene expression profiles, as many other omics approaches, yield noisy data and more robust approaches to the analysis of gene expression data should be explored.

Here we present a novel method that, instead of characterizing a perturbation signature by listing the significantly up-regulated and down-regulated genes, uses these data to estimate the activation of transcription factors (TFs). Several online resources integrate experimental datasets and/or computational approaches to compile lists of genes whose expression is modulated by a TF [13]. This information permits to convert a gene expression profile into a Transcription Factor Activation Profile (TFAP) by estimating TF activities from the expression of their target genes. It is then possible to infer the impact of a drug on activation of a TF by looking at the fraction of target genes that are up- or down-regulated after drug treatment. This procedure is often referred to as TF enrichment analysis. We show that this approach is less sensitive to experimental noise when compared with conventional expression profile methods and we used it to infer drugs that are likely to induce myeloid differentiation of HL-60, a cell line derived from a patient with acute myeloid leukaemia.

## 2. Results

### 2.1 Identification of compounds promoting differentiation in Acute Myeloid Leukaemia by TFAP approach

Repositioning strategies based on comparison of transcriptional signatures have encountered some success [7]. However, they suffer from experimental noise, which characterizes high throughput experiments. In addition, transcriptional signatures are largely cell type specific and profiles derived from experiments in one cell type are often not transferable to different cell types.

We reasoned that comparison of drug profiles, defined as lists of transcription factor (TF) activation levels, rather than transcriptional profiles, could offer a more robust strategy. TF activation can be deduced from a large number of measurements of the different target genes and as such it is less sensitive to the experimental variability of each single gene-expression data point. In this strategy (**Figure 1**) activations of TFs are not derived from the levels of their messenger RNAs (mRNAs), as mechanisms other than modulation of transcription are in many cases at the basis of their activation. TF activations are rather deduced from the differential expression of the mRNAs of their target genes (see Material and Methods). Target genes can be associated to TFs by a variety of criteria. These include literature-curation, chip-seq, co-expression and *in silico* prediction of TF binding sites [13], [14]. To build drug-TF activation signatures we have used the ChEA3 resource (https://amp.pharm.mssm.edu/chea3/). ChEA3 accepts as input a list of significantly up- and down-regulated genes in a given condition, it confronts this list with lists of genes that are annotated to each TF and returns a table containing all the transcription factors whose annotated target genes are enriched in the input gene list.

**Figure 1.**
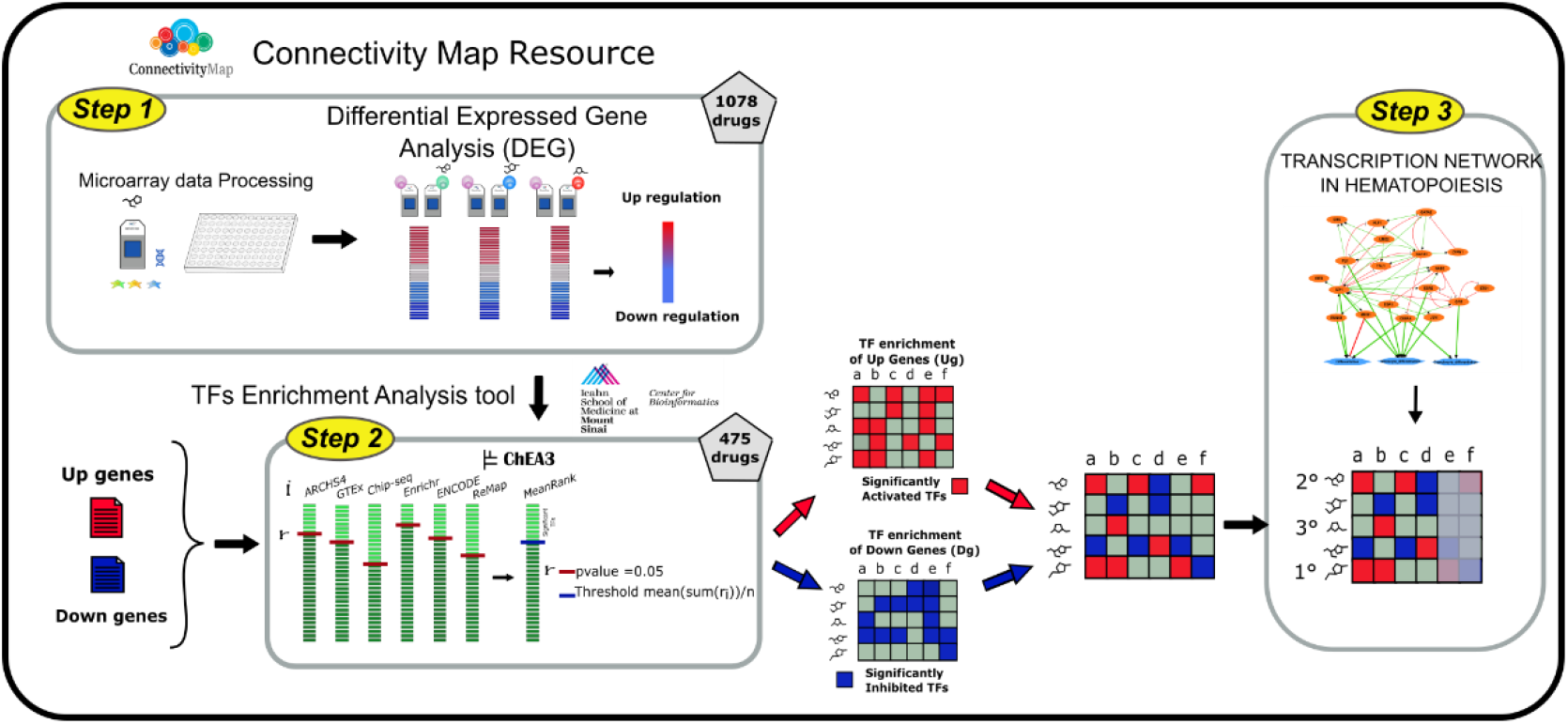
Workflow of the proposed strategy. **Step 1** Drug-specific gene expression profiles are derived from the Connectivity Map resource and compile lists of up- and down-regulated genes are compiled. **Step 2**: Transcription factor (TFs) Enrichment analysis with the ChEA3 tool. The results are represented as two matrices (Drugs x TFs) where significantly activated and inactivated TFs are shown with a red and blue background respectively. The two matrices are integrated; the TFs whose targets are enriched in both gene lists are not considered and labelled with a grey background in the matrix. **Step 3**: the matrix was simplified by focusing on those TFs that are involved in the biological function of interest, in our case monocyte and granulocyte differentiation.

To test the premise that TF activation profiles are effective in filtering noise we first devised a pilot study. The rational of the test is schematically illustrated in **Figure 2** where we have represented as two-dimensional tSNE maps [15] the multi-dimensional dataset represented either as feature vectors of gene expression levels or TF activation. We applied our analysis to the CMap dataset, which consists of transcription profiles after perturbation by approximately 1078 drugs in three different tumor cell lines. The gene expression profiles of the different cell lines are rather different and the response to the same drug needs not be necessarily similar as it is likely to have some cell-type specificity [16]. Thus, in a two-dimensional tSNE map representation of the multiparametric dataset, expression profiles tend to cluster according to cell type rather than according to drug (**Figure 2A**). On the other hand, the conversion of the transcriptional profiles into TF activation profiles is surmised to make the perturbations by the same drug on the different cell lines more similar, at least for some drugs, as a drug targets the same transcriptional circuit in all cell types (**Figure 2B**). To test this hypothesis we measured, in the two-dimensional tSNE map, the distance between the points corresponding to the same drug perturbation (same data point-symbol in **Figure 2 A, B**). This measure is taken as a proxy of profile similarity. The expected and observed distance distributions are shown **Figure 2C** and **2D** respectively. In **Figure 2D** the observed distance distributions obtained by applying the gene expression (top) and TF activation profiles (lower plot) are compared with the distributions obtained by performing the same analysis on randomized transcription and TF profiles. In both cases the observed distributions are significantly different from the one based on randomized transcription profiles, the observed one being more shifted toward short distances as would be expected if many drugs would produce similar perturbations of the transcriptional profiles of the different cell lines. On the other hand, the distribution curves of the analysis using as features TF activation (lower plot), are evidently more different. This observation is quantitatively confirmed by a Kolmogorov-Smirnov test (**Figure 2E**) supporting the notion that the approach based on the inference of TF activation is less sensitive to data noise as it is capable to recognize similar effects in the perturbations of different cell lines with the same drug, in a larger number of instances.

**Figure 2.**
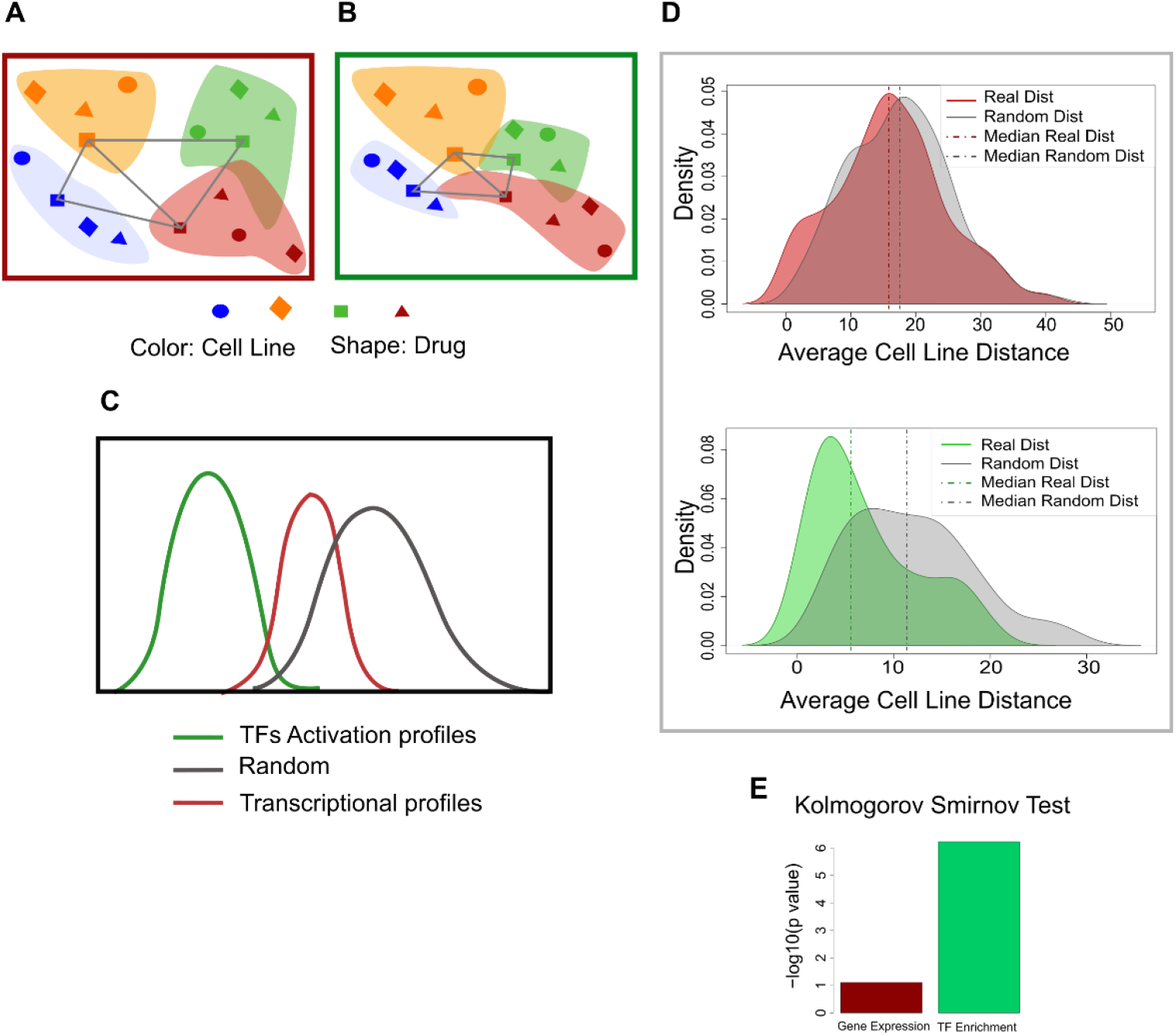
Comparison of the transcription-profile perturbations caused by different drugs in different cell lines. **(A)** Schematic representation of bidimensional tSNE maps of the multidimensional transcriptional profiles of different cell lines are incubated with different drugs. Cell lines are represented with different colors while different drugs have different shapes. **(B)** Same as A after the transformation of the expression profiles into TF activation profiles. **(C)** Inferred distribution of distances between profiles of different cell lines incubated with the same drug. **(D)** Experimental distribution of tSNE distances when profiles are defined as gene expression vectors (top) or TF activation vectors(lower plot) The distance distributions observed in randomized datasets are shown with a gray background. **(E)** Kolmogorov-Smirnov test of the significance of the differences between profile distances in experimental and random datasets.

### 2.2 Identification of drugs that activate master regulators of myeloid differentiation

To prioritize drugs for testing their ability to promote myeloid differentiation we next applied the newly developed strategy to the HL-60 CMap dataset. HL-60 is relevant for our goal of identifying drugs that promote differentiation in AML as it is a human leukaemia cell line [17] that can be induced to differentiate into monocytes and granulocytes. We first aimed at listing significantly up- and down-regulated genes when HL-60 cells are treated with 1078 different drugs in the CMap dataset. Only for 475 of these perturbation experiments it was possible to compile lists of significantly modulated genes. These lists were used as input in the ChEA3 resource to identify transcription factors that are likely to be up- or down-regulated by each drug. This approach yielded a matrix of 475 compounds times 1632 TFs, where drug-activated and inactivated TFs are labelled in red and blue background respectively in **Figure 3A**. Next, we screened the literature searching for reports of experimental evidence about the regulatory circuits involved in haematopoiesis. The different differentiation steps are regulated by a complex interplay among numerous transcription factors [18]. We focused on 7 of these that either play a major role as key regulators of haematopoietic stem cells (HSCs) differentiation and/or are activated in the terminal steps controlling granulocyte and monocyte differentiation (**Supplementary Table 1**).

**Figure 3.**
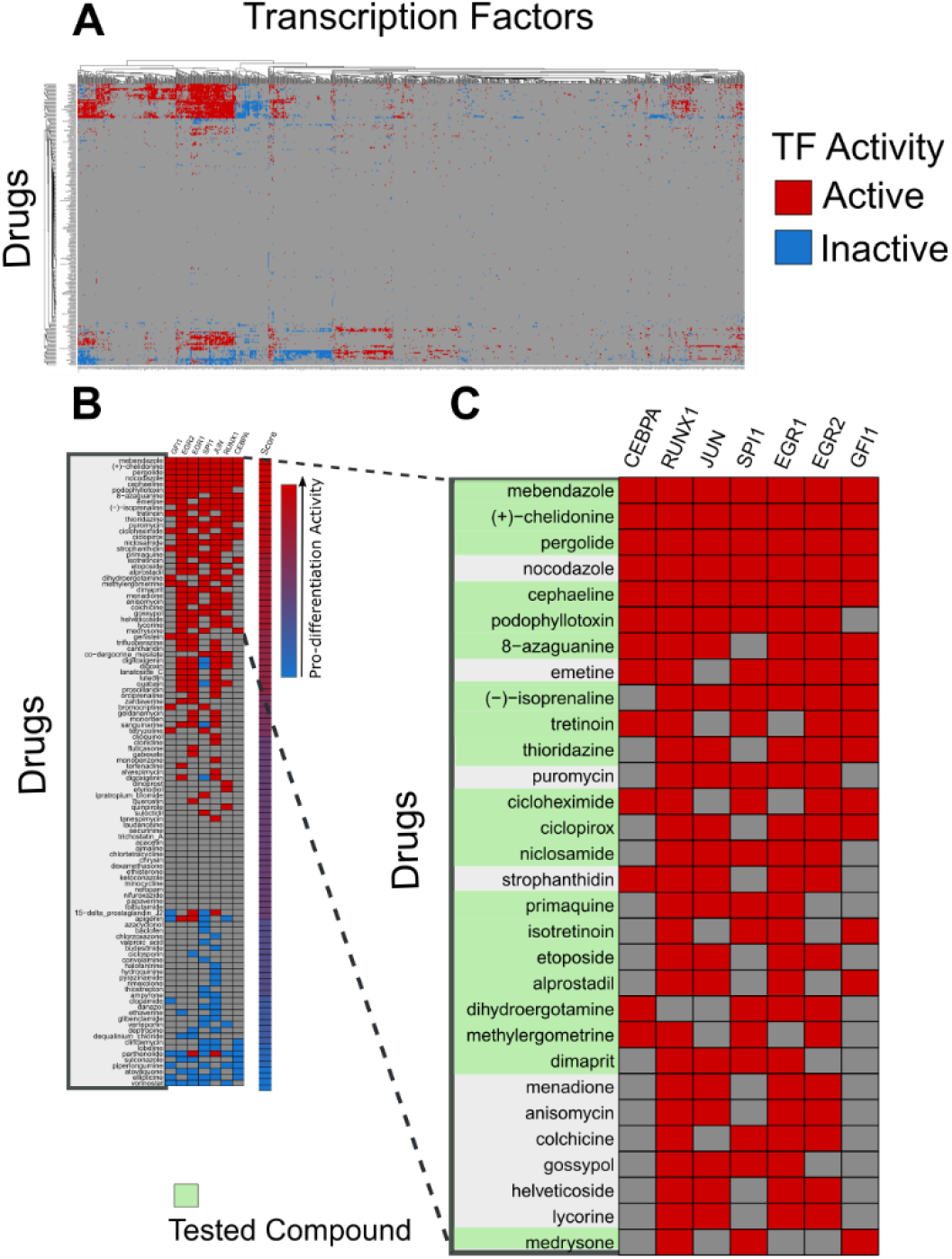
Transcription factors activated or inhibited by drugs in HL-60. **(A)** Matrix reporting the results of the ChEA3 analysis of genes up- and down-regulated by incubation of HL-60 cells with 475 drugs. Each row corresponds to a drug while transcription factors are in columns. A red background denotes that the transcription factor in the column is activated by the chemical in the row. Conversely a blue background indicates inactivation. **(B)** The matrix in A was simplified by maintaining only the results related to the seven transcription factors that promote monocyte and granulocyte differentiation. Only the data of the 107 chemicals that have an effect on transcription factors involved in haematopoiesis are shown. Chemicals were ordered according to a “pro-differentiation score” computed by adding up, for each chemical, the number of activated transcription factors and subtracting the number of the inhibited ones. **(C)** Close up of the top-ranking pro-differentiation compounds. The chemicals that have been tested experimentally have a light green background.

In order to identify the drugs that affect the activity of these seven master regulators of myeloid differentiation, we first simplified the matrix that links drugs to transcription factor activation and only considered the 107 chemicals that, according to the ChEA3 resource have an effect on the seven TFs involved in myeloid differentiation. The resulting reduced matrix is shown **Figure 3B** where we have ordered the compounds according to the number of pro-differentiation TFs that they activate. The ranking method is described in more detail in the Materials and Methods section. In **Figure 3C** we have shown an enlargement of the section of the matrix listing the top 30 compounds in the rank list. Among the 107 selected compounds, we notice, in the high-ranking positions, tretinoin, a known differentiation inducer of leukemic cells [19] thus providing confidence in the potential of the approach.

### 2.3 Experimental validation of drugs inferred to induce granulocytic differentiation

To validate our strategy, we set out to test some of the inferred pro-differentiation drugs by evaluating their potential to induce differentiation of HL-60. To this end we tested 22 of the top-ranking compounds highlighted in in **Figure 3C**. HL-60 were treated with the selected drugs for four days (**Figure 4A**) and granulocyte differentiation was assessed by the nitroblue tetrazolium (NBT) assay. In **Figure 4B** we have graphically summarized these results by labelling in the matrix of **Figure 3C** the compounds that did or did not show a pro-differentiation effect with orange and yellow backgrounds respectively. Interestingly, 10 of the 22 drugs significantly induced granulocytic differentiation in HL-60 as shown by NBT quantitation and staining (**Figure 4C and E**). Quantitation showed that, in addition to tretinoin, our positive control, nine additional drugs, namely mebendazole, etoposide, 8-azaguanine, alprostadil, dimaprit, dihydroergotamine, methylergometrine and pergolide were able to induce significant differentiation in HL-60 (**Figure 4C**). After treatment, viability was measured by the trypan blue exclusion test to evaluate drug toxicity in these conditions. Three of the 22 compounds showed a toxic effect as fewer than 20% of the cell survived the treatment. These compounds were not characterized further (**Figure 4D**). To further verify the robustness of the computational strategy, we randomly chose 14 drugs (**Supplementary Figure 2B**) from the 1078 drugs considered in the computational screening and we tested them for toxicity and for the ability to induce granulocytic differentiation in HL-60 by the trypan blue exclusion test (**Supplementary Figure 2E**) and the NBT assays (**Figure 4F, Supplementary Figure 2D**). No such drug showed significant toxicity. Similarly, none of the randomly selected drugs was able to induce HL-60 differentiation into granulocytes. This result was confirmed by quantitation of the NBT assay (**Figure 4F**). Thus, randomly chosen drugs, differently from those inferred by the computational approach have negligible probability to induce a significant differentiation of HL-60 (**Figure 4G**). We conclude that this approach, being sensitive and specific is suitable for inferring drugs able to induce granulocytic differentiation.

**Figure 4.**
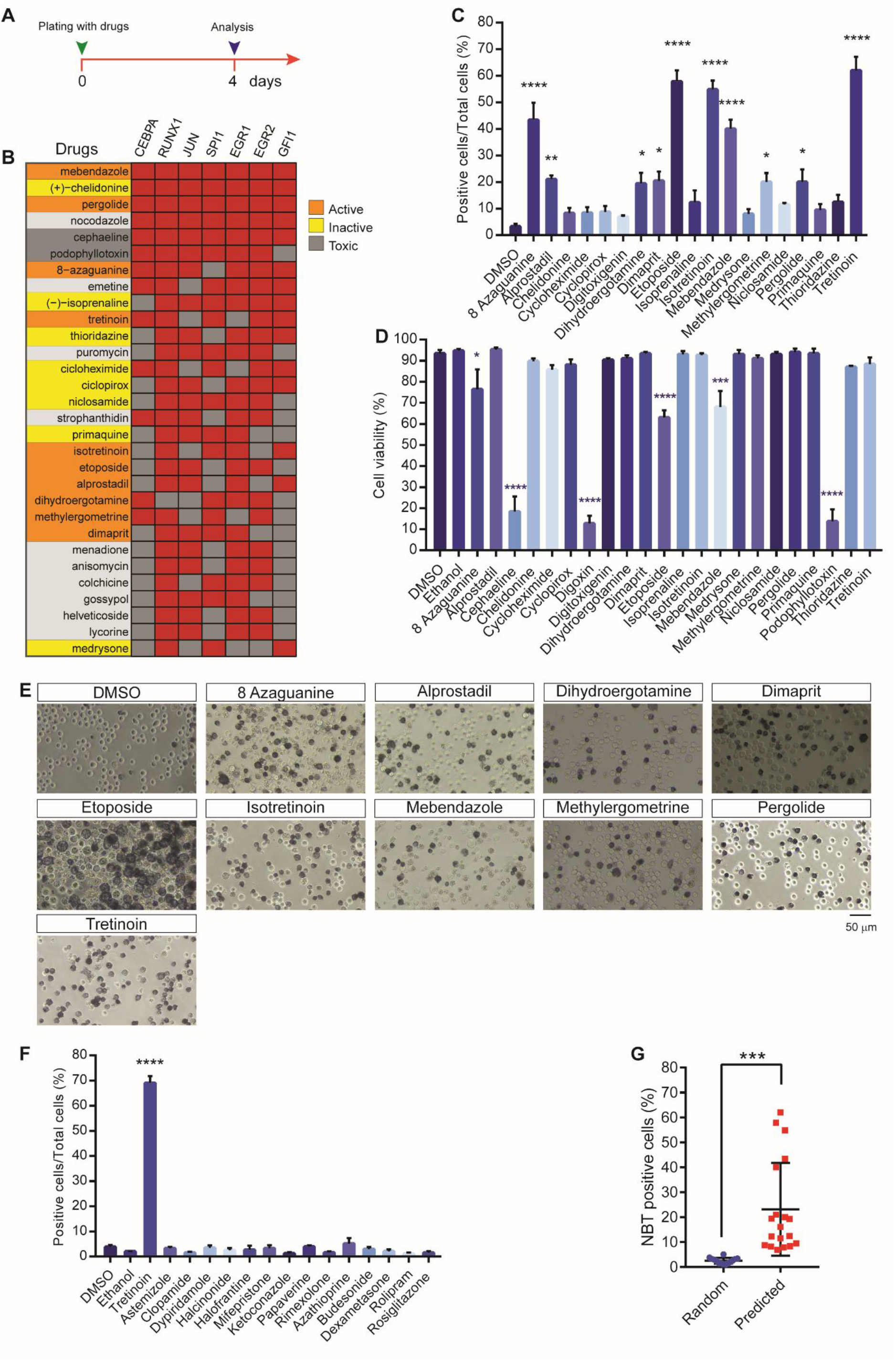
Experimental validation of inferred pro-differentiation and randomly selected drugs in HL-60. **(A)** Time line of the experiment to test the pro-differentiation activity of drugs on HL-60 cells. **(B)** Top 30 ranking compounds with corresponding experimental outcomes. Compounds are ranked according to their “pro-differentiation score”. Compounds that have shown or not a significant activity on HL-60 differentiation are labelled with and orange or yellow background respectively. A compound that was toxic at the concentrations used in the assay is labelled with a grey background. **(C)** Quantitation of NBT assay of HL-60 treated with the drugs listed in **Supplementary Figure 2A**. All drug treatments were carried out for four days. The concentration for each drug is reported in Table 1 in Materials and Methods section. The NBT assay results were plotted as percentage of positive cells over the total cells. **(D)** Cell viability after four days of selected drug treatment. Cell viability was assessed by the Trypan blue exclusion test. **(E)** Representative light microscope images of NBT staining in HL-60 after four days of drug treatment. Scale bar is 50 μm. **(F)** Quantitation of the NBT assay of HL-60 treated with the randomly selected drugs listed in Supplementary Figure 2B. The concentration for each drug is reported in in Table 1 in the Materials and Methods section The NBT assay results were plotted as percentage of positive cells on the total cell number. **(G)** Dot plots of the results of NBT assays on HL-60 treated with randomly selected compounds (blue circles) or predicted compounds (red squares). Each circle or square represent the mean of the percentage of NBT positive cells over total cells of three biological replicates. Statistical significance was evaluated using a Student t-test (n = 3). Data are presented as mean ± SEM. * p ≤ 0.05, ** p ≤ 0.01, *** p ≤ 0.001Data in **(C)**, **(D)** and **(F)** are represented as means of three biological replicates (n = 3) ± SEM. Statistical analysis was performed using One-way ANOVA. Significance * p ≤ 0.05, ** p ≤ 0.01, *** p ≤ 0.001, **** p ≤ 0.0001 and are related.

### 2.4 Drugs that are high in the ranking list increase the levels of surface markers expressed by differentiating HL-60 cells

The increase in the expression of specific cell surface markers correlates with the differentiation of HL-60 into mature granulocytes. Thus, we measured the levels of the integrin alpha M (CD11b), which is expressed in mature granulocytes and regulates cell adhesion and migration [20]. We monitored CD11b expression by flow cytometry after four days of treatment with the compounds that scored positive in the NBT assay. The histograms in **Figure 5A** reports the flow cytometric analysis of CD11b expression in HL-60 treated with the different drugs. We observe that 7 of the 10 compounds significantly express CD11b thus confirming their ability to promote cell differentiation (**Figure 5B**). We also monitored the expression of the transferrin receptor protein 1 (TfR1), an integral membrane protein expressed in proliferating cells also known as CD71 [21], [22]. Moreover, we evaluated the levels of the cyclic ADP ribose hydrolase (CD38) a glycoprotein which is expressed by HL-60 only when cells are treated with retinoids [23] (**Figure 5C**). Not all the pro-differentiation compounds induce expression of CD38, notwithstanding their ability to induce differentiation in HL-60 (**Figure 5D**). Of note, the compounds that induce more efficiently differentiation, cause a decrease of CD71 expression (**Figure 5E**) in accord with the notion that differentiated cells loose proliferation potential [24]. We also looked at the expression of Carcinoembryonic antigen-related cell adhesion molecule 8 (CD66b), a cell adhesion molecule that is expressed by granulocytes [25]. Most samples treated with the compounds contain cells expressing this marker, albeit at low levels (**Figure 5F**) and only three compounds promote a significant increase in CD66b expressing cells (**Figure 5G**). This is not unexpected as in mature granulocytes the expression of CD66b is low and combined treatments with different stimuli are necessary for a full expression of this marker [26]. We conclude that drugs that are rated high by our bioinformatic approach promote the expression of cell surface markers that characterize granulocytes.

**Figure 5.**
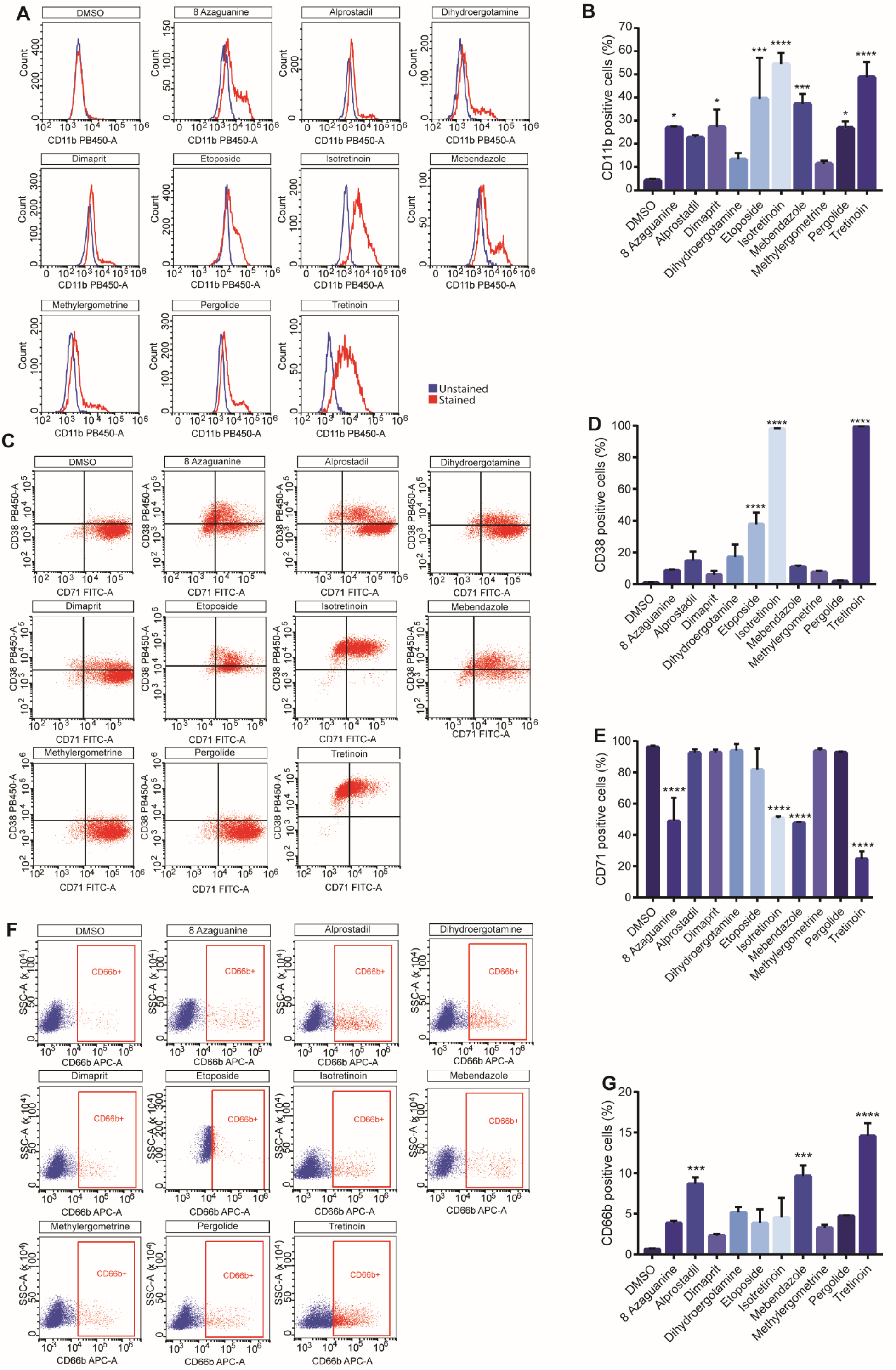
Expression of differentiation markers in HL-60. **(A)** Representative histogram of flow cytometric analysis of CD11b expression in HL-60 after four days of treatment with the indicated drugs. **(B)** Quantification of CD11b expression in HL-60 after four days of treatment. **(C)** Scatter dot plots representing HL-60 population analysed by flow cytometry with anti CD71 and anti CD38 fluorescent antibodies. The representative dot plots show the percentage of CD71 and CD38 positive and negative cells after four days of treatment with different drugs. **(D)** Quantification of CD38 expression in HL-60 after four days of treatment of three different biological replicates. **(E)** Quantification of CD71 expression in HL-60 after four days of treatment of three different biological replicates. **(F)** Representative histogram of flow cytometric analysis of CD66b expression in HL-60 after four days of treatment. Red gate encloses CD66b positive cells. **(G)** Quantification of CD66b expression in HL-60 after four days of treatment. Statistical analysis was performed using ordinary One-way ANOVA. Data are presented as mean ± SEM. *** p ≤ 0.001, **** p ≤ 0.0001. All the drug treatments were carried out for four days. The concentration for each drug is reported in **Table 1** in Materials and Methods section.

### 2.5 Drugs that induce a DNA damage response also promote a differentiation program in HL-60

By going over the literature reporting cell mechanisms triggered by the compounds that we have observed to promote cell differentiation, we noticed that some are known inducer of the DNA damage response. Among them, i) 8-azaguanine, a purine analogue that incorporates into ribonucleic acids and interferes with physiological biosynthetic pathways [27], ii) mebendazole that binds to the colchicine-binding domain of tubulin thereby inhibiting its polymerization [28] and iii) etoposide that triggers the DNA damage response by inhibiting topoisomerase II [29]. We set out to further investigate the potential causal link between DNA damage response and myeloid differentiation. We know from experiments in our group that idoxuridine induces DNA damage by incorporating into the DNA. The DNA damage in turn promotes osteogenic differentiation of mesangioblasts [30]. Hence, we tested idoxuridine in our model system to strengthen the hypothesis of a link between DNA damage and differentiation. After 4 days of treatment with 10 μM idoxuridine the number of cells that stain blue in the NBT assay increases significantly (**Figure 6A and B**). Moreover, idoxuridine treatment leads to a marked increase in CD11b and CD66b expression (**Figure 6C, H**) and a decrease in CD71 expression (**Figure 6E-G**) while, as observed earlier for 8-azaguanine and mebendazole, it does not induce the expression of CD38 (**Figure 6E-F**), suggesting a different differentiation mechanism when compared to tretinoin. These results, taken together, confirm a causal link between the DNA damage response and differentiation of leukaemic stem cells.

**Figure 6.**
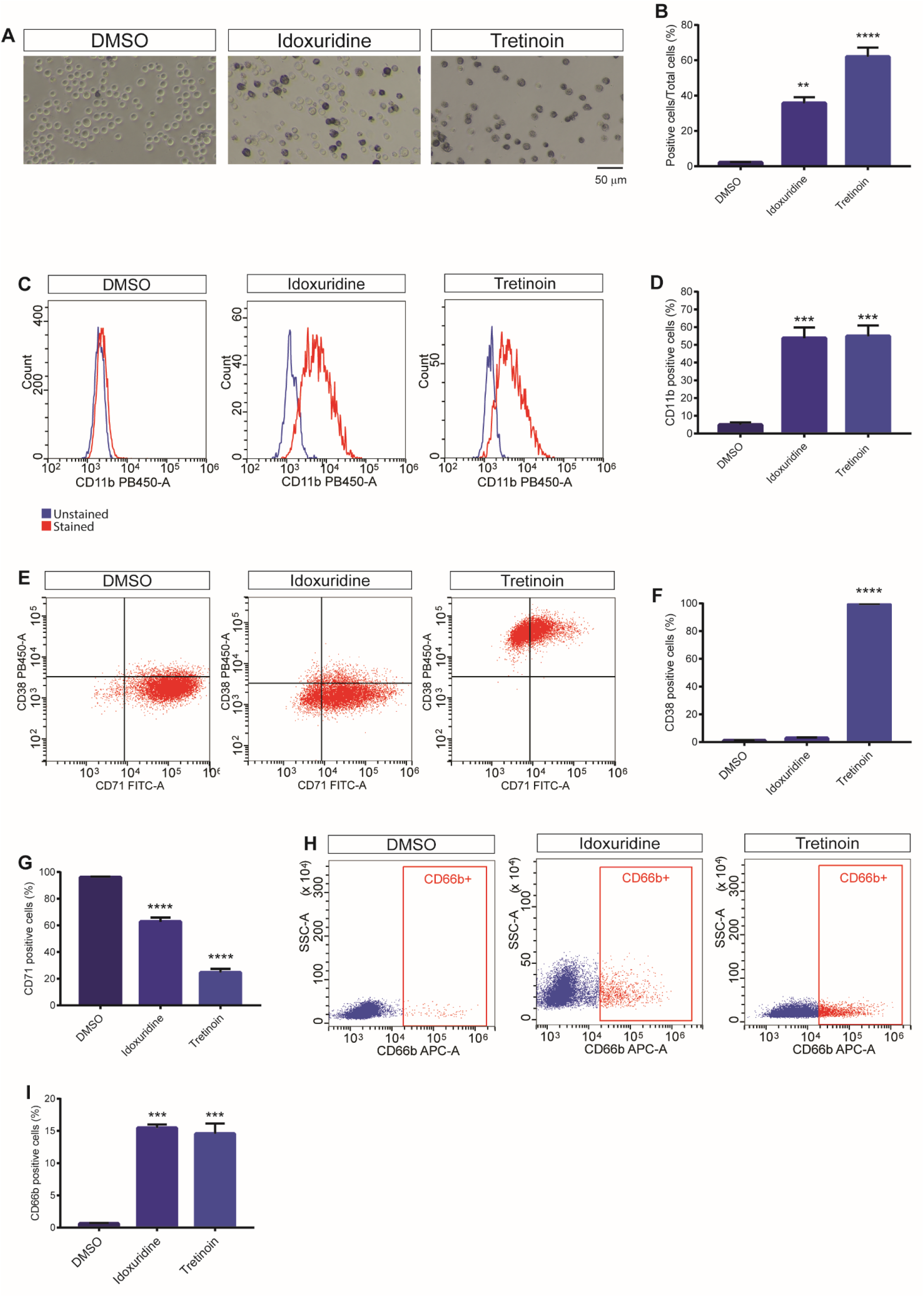
Idoxuridine induces differentiation of HL-60. **(A)** Representative light microscope images of NBT staining of HL-60 after four days of treatment with idoxuridine (10 μM) and Tretinoin (1 μM). The scale bar is 50 μm. **(B**) Quantitation of the assay in A. **(C)** Intensity distribution and quantitation **(D)** of flow cytometric analysis of CD11b expression in HL-60 treated for four days with idoxuridine (10 μM) and tretinoin (1 μM). **(E)** Scatter dot plots and quantitation of the expression of CD38 **(F)** and CD71 **(G)** after idoxuridine and tretinoin treatment. **(H)** Representative dot plots of flow cytometric analysis and quantitation **(I)** of CD66b expression in HL-60 treated for four days with idoxuridine and Tretinoin (1 μM). The red gate represents CD66b positive cells. Statistical analysis was performed using ordinary One-way ANOVA. Data are presented as mean ± SEM. ** p ≤ 0.01, *** p ≤ 0.001, **** p ≤ 0.0001. All the drug treatments were carried out for four days in biological triplicates. Drug concentrations are reported in **Table 1** in Materials and Methods section.

### 2.6 The selected drugs are also efficient in inducing differentiation in a non-promyelocytic cell line

One of the limits of the drugs inducing leukaemic cell differentiation is that they are not equally efficient in inducing differentiation of all AML cell types. For this reason, we tested the potential of some of the drugs that we have identified to trigger differentiation in a second cell type. To this end we chose THP1, a human monocytic cell line isolated from the peripheral blood of a one-year-old male patient suffering from acute monocytic leukaemia [31]. This cell line was classified as FAB M5 subtype and can differentiate into macrophage like cells [32]. We performed the NBT assay in THP1 cells after four days of treatment with the drugs (**Figure 7A)**. While no drug showed significant toxicity (**Figure 7B**), they induced macrophage differentiation in the THP1 cell line as shown by NBT staining (**Figure 7C**). These analyses demonstrated the potential of the newly identified pro-differentiation drugs to induce differentiation of, at least, two human leukaemia cell types.

**Figure 7.**
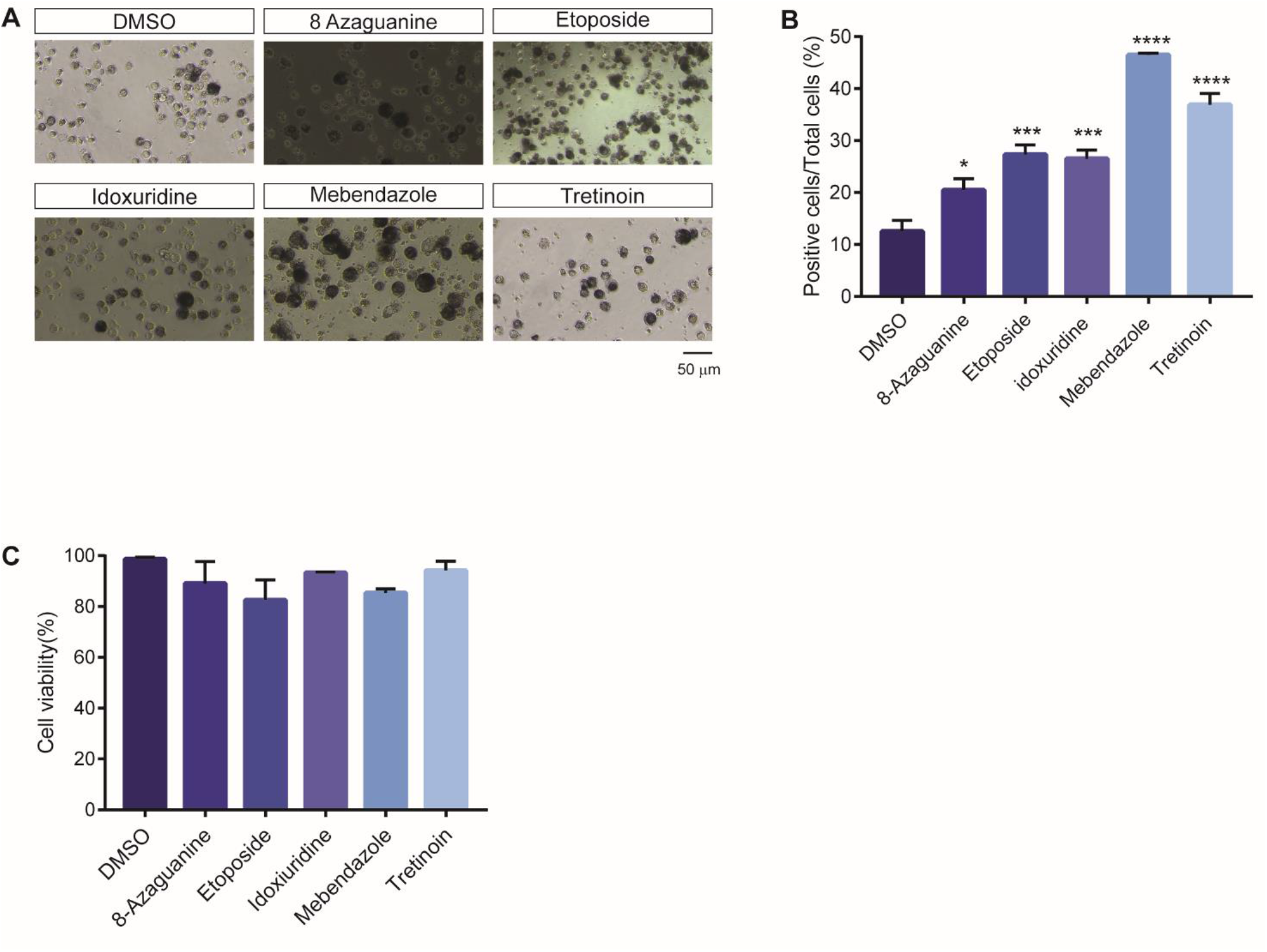
NBT assay of THP1 cell line differentiation. **(A)** Representative light microscope images of NBT staining of THP1 after four days treatment with the drugs. The scale bar is 50 μm. **(B)** Bar plot reporting THP1 cell viability after four days of treatment with the drugs. Cell viability was assessed by the Trypan blue exclusion test. Data points are averages of three biological replicates. **(C)** Quantitation of the NBT assay on drug treated THP1 samples. All drug treatments were carried out for four days. The concentration of each drug is reported in Supplementary Table 1. The NBT assay results were plotted as percentage of positive cells over the total cells. Statistical analysis was performed using One-way ANOVA. Data are presented as mean ± SEM. of three biological replicates. Significance * p ≤ 0.05, *** p ≤ 0.001, **** p ≤ 0.0001 and are related.

## 3. Discussion

Drug repurposing has attracted increasing attention in recent years [33]–[36]. Over the past decades several drugs have been repurposed for new therapeutic uses, e. g. sildenafil, minoxidil, aspirin, valproic acid, [33], [37]–[39]. In this context, we developed a new strategy to screen drugs *in silico* looking for compounds that could be candidates for differentiation therapy in Acute Myeloid Leukaemia (AML). AML is a heterogeneous and aggressive disease with poor survival rate [40]. Its main characteristic is the inability of haematopoietic stem cells to terminally differentiate accompanied by an increase of their self-renewal potential [41]. Current therapy approaches, based on chemotherapy, albeit somewhat aggressive, do not completely revert the adverse pathological outcome in AML patients. These considerations stimulated alternative approaches such as for instance differentiation therapy. The efficacy of All-Trans Retinoic Acid (ATRA) as differentiation agent in AML subgroups treatment [19] underscores the potential of this approach [42] and motivates the identification of additional compounds that promote the differentiation of leukemic stem cells. To this end we set out to develop a novel computational approach in order to relocate compounds with the potential to reprogram leukemic stem cells. By interrogating the Connectivity Map (CMap), we performed a differential gene expression analysis of the human leukaemia cell line HL-60, treated with several drugs. We focused on compounds that enhance the expression of genes regulated by the transcription factors that are known to play a crucial role in myeloid differentiation. Because of the interconnected nature of cell signaling, targeting transcription factors, which act downstream in the signaling process, is a more specific strategy than focusing on the expression on upstream cell-signaling nodes. The strategy that we have developed allows to overcome some of the limitations of standard drug repurposing approaches based on the screening of a large number of compounds by cell assays [43], [44]. Our strategy, however, has some limitations as the CMap resource has incomplete drug coverage. In addition, all the data were obtained by using a microarray platform, a technology that is by now obsolete [45]. In spite of this, the method has proven to have value as it provided us with a list of potential pro-differentiation compounds that include drugs already used in differentiation therapy. This gave us confidence in the potential of our computational repositioning method. In order to validate the effect of candidate pro-differentiation drugs, we performed differentiation assays *in vitro*. The observation that 10 of the 22 inferred drugs, could promote granulocytic differentiation of HL-60, supported the value of the computational method and estimated to approximately 50% the frequency of false positives. Moreover, to evaluate the frequency of false negatives, we randomly selected 14 drugs that had a low position in the ranking list. None of them showed a significant potential to stimulate differentiation thus establishing a higher limit of approximately 10% in the frequency of false negatives. Among the pro-differentiation compounds, we identified mebendazole, an anti-helminthic agent recently proposed by Yulin Li and colleagues as a drug for AML differentiation therapy [46]. The authors elaborated a computational approach, referred as Lineage Maturation Index (LMI), to define the changes in differentiation state of haematopoietic malignancies based on their gene expression profiles. This approach similarly to ours also used the CMap dataset and compared transcriptional profile instead of transcription factor activation profiles. The approach produced a list of putative drugs that were subsequently tested *in vitro*. Despite being the two approaches based on the same experimental dataset, we consider our computational strategy more efficient and sharper in identifying pro-differentiative compounds. Indeed, by comparing the list of the pro-differentiative drugs obtained by the two approaches, the strategy proposed here identified additional compounds not recognized by the LMI approach. These include 8-azaguanine, alprostadil and methylergometrine. This underlines the power of our strategy that is, in addition, highly versatile as it can be relatively easily adapted to different biological processes and drug repurposing projects. The list of new pro-differentiation drugs allowed us to contribute to shed some light on the mechanisms governing differentiation in myeloid cells. In particular, we observed that drugs that induce differentiation more efficiently also trigger a DNA damage response. The DNA damage response has already been discussed as a target to modulate differentiation of myeloid leukaemia cells [47]–[49]. This consideration prompted us to further investigate the correlation between DNA damage and myeloid differentiation. To this end, we tested idoxuridine (IdU), a molecule that promotes cell differentiation by triggering DNA damage response in a completely different cell type and experimental set up [30]. We observed that IdU induces myeloid differentiation with a potency that is comparable to that of ATRA. More effort will be necessary to understand the details of the molecular mechanisms underlying IdU pro-differentiation activity. Possibly IdU triggers HL-60 granulocytic differentiation via a mechanism that is different from that of ATRA, as shown by the different expression of CD38 stimulated by the two drugs. Several lines of evidence suggest that CD38, a type II transmembrane ectoenzyme and receptor, may be a key driver of ATRA induced differentiation given that, after ATRA treatment, HL-60 show a dramatic upregulation of CD38 [50], [51]. We observed that, by contrast, IdU does not induce an increase in CD38 expression in HL60, despite being able to promote granulocytic differentiation, suggesting a different mechanism of action [52]. Further investigations in primary human AML cells may be helpful to assess the potential of the selected drugs for use in differentiation therapy of AML, possibly in combination with already approved drugs. In conclusion, this work further emphasizes the potential of repurposing strategies and the contribution that computational approaches provide in accelerating the addition of new items to the therapeutic toolbox.

## 4. Materials and Methods

### 4.1 Connectivity Map

The Connectivity map is a resource (CMap build 02) [12] collecting 6100 gene expression profiles of 5 human cell lines treated with 1309 Food and Drug Administration (FDA) approved compounds. One of the cell lines used in this project is HL60, a human leukaemia cell line whose expression profiles was determined after treatment with 1078 compounds. However, the number of drug treatments and control samples are not always compatible with statistical analysis. As a consequence, we were only able to determine a significant perturbation of the transcriptional signature for 475 compounds. The expression data were processed and normalized. Differential gene expression analysis was performed by using the R *Limma* package [53]. For each drug treatment this procedure generated two lists of significantly up- and down-regulated genes, which were used as input in the TF enrichment tool ChEA3.

### 4.2 ChEA3 and integration of TF enrichment results

The analysis of the transcription factors whose target genes are enriched in the list of genes that are up/down regulated after drug treatment was obtained by the ChEA3 tool (https://amp.pharm.mssm.edu/chea3/). To estimate the TF target enrichment ChEA3 uses seven different resources (ARCHS4 Coexpression, ENCODE, ChIP-seq, EnrichrQueries, GTExCoexpression, Literature, ReMapChIP-seq) each using a different method to compile list of genes that are controlled by each transcription factor. Each resource uses different methods to compile list of genes that are controlled by each transcription factor. For each chemical perturbation, they return a list of activated transcription factors ranked according to the p-value of the enrichment of its target genes in the lists of up/down regulated genes under perturbation conditions. We used the *MeanRank* method to combine the results of enrichment analyses in the seven different resources. For each TF the *MeanRank* method computes the mean of the rank positions in the output lists of the seven resources. This integration strategy was shown to perform better than each single method (Kennan et al 2019).

The MeanRank, however, does not allow to define a p-value for a specific position in the ranked list. We thought of an empirical method for establishing a threshold position above which a TF could be considered significantly “activated”. To this end for each ranking list we noted the position of the transcription factor whose target-enrichment p-value was just below the significance of 0.05. We next computed the average of the threshold positions in the seven resources. Finally, the TF with an average position higher than this average threshold position were deemed “significantly activated”. At the end of this process for each chemical we have a list of transcription factors whose targets are significantly more numerous in the list of up- or down-regulated genes as evaluated by the annotations in the seven different resources.

### 4.3 TF filtering and drug ranking procedure

The drugs that are candidates for induction of myeloid differentiation are those that activate the TFs that are necessary for differentiation while inactivating fewer of them. Only transcription factors specifically involved in monocyte and granulocyte differentiation (MGD) were considered. In particular, we focused on those seven transcription factors that promote granulocyte or monocyte differentiation. To obtain a ranking list of pro-differentiation compounds, for each drug we counted the number of myeloid differentiation TFs involved in myeloid differentiation that are significantly activated and subtracted those whose targets are enriched in the list of down-regulated genes. This score was dubbed “pro-differentiation” score.

### 4.4 Cell cultures

HL-60 were purchased from ATCC (American Type Culture Collection) (#CCL-240™) and were cultured at density of 5.0 × 10^5^ cells/mL in T-75 cm^2^ flasks (Corning^®^, #431464U) using growth medium consists of Iscove’s Modified Dulbecco’s Medium (IMDM) (ATCC^®^ 30-2005™) supplemented with 20% v/v heat inactivated Fetal Bovine Serum (FBS) (Euroclone, #ECS0180L), 1 mM sodium pyruvate (Sigma-Aldrich, #S8636), 10 mM 4-(2-hydroxyethyl)-1-piperazineethanesulfonic acid (HEPES) (Sigma, #H0887) and 100 U/ml penicillin/100 μg/ml streptomycin (Thermo Fisher Scientific, #15140122) at 37°C in 5% CO_2_ atmosphere. Cultures were maintained by the addition of fresh medium or replacement of medium every 2-3 days. Cell concentration did not exceed 1 × 10^6^ cells/mL.

THP1 were purchased from ATCC (American Type Culture Collection) (#TIB-202™) and were cultivated at a density of 2.0 × 10^5^ cells/mL in T-75cm^2^ flasks (Corning^®^, #431464U) in Roswell Park Memorial Institute (RPMI) 1640 medium (Gibco, #11875093) supplemented with 10% v/v heat inactivated Fetal Bovine Serum (FBS) (Euroclone, #ECS0180L), 1 mM sodium pyruvate (Sigma-Aldrich, #S8636), 10 mM 4-(2-hydroxyethyl)-1-piperazineethanesulfonic acid (HEPES) (Sigma, #H0887) and 100 U/ml penicillin/100 μg/ml streptomycin (Thermo Fisher Scientific, #15140122 at 37°C in 5% CO_2_ atmosphere. Cultures were maintained by the addition of fresh medium or replacement of medium every 3-4 days. Cell concentration did not exceed 8 × 10^5^ cells/mL.

### 4.5 Drug treatments

HL-60 were treated with different drugs according to Gupta Et al. [54]. Cells were diluted to 3 × 10^5^ cells/ml in IMDM growth medium and incubated overnight to obtain a population of exponentially growing cells. The following day cells were centrifuged at 300 × g for 7 minutes at room temperature and resuspended at the density of 3.0 × 10^5^ cells/mL in IMDM growth medium supplemented with the different drugs. Chemicals were added to the growth medium at the same concentrations used in the connectivity map protocol. The drug concentrations used for each compound are shown in **Table 1** Drug treatment was carried out for 4 days.

**Table 1:**
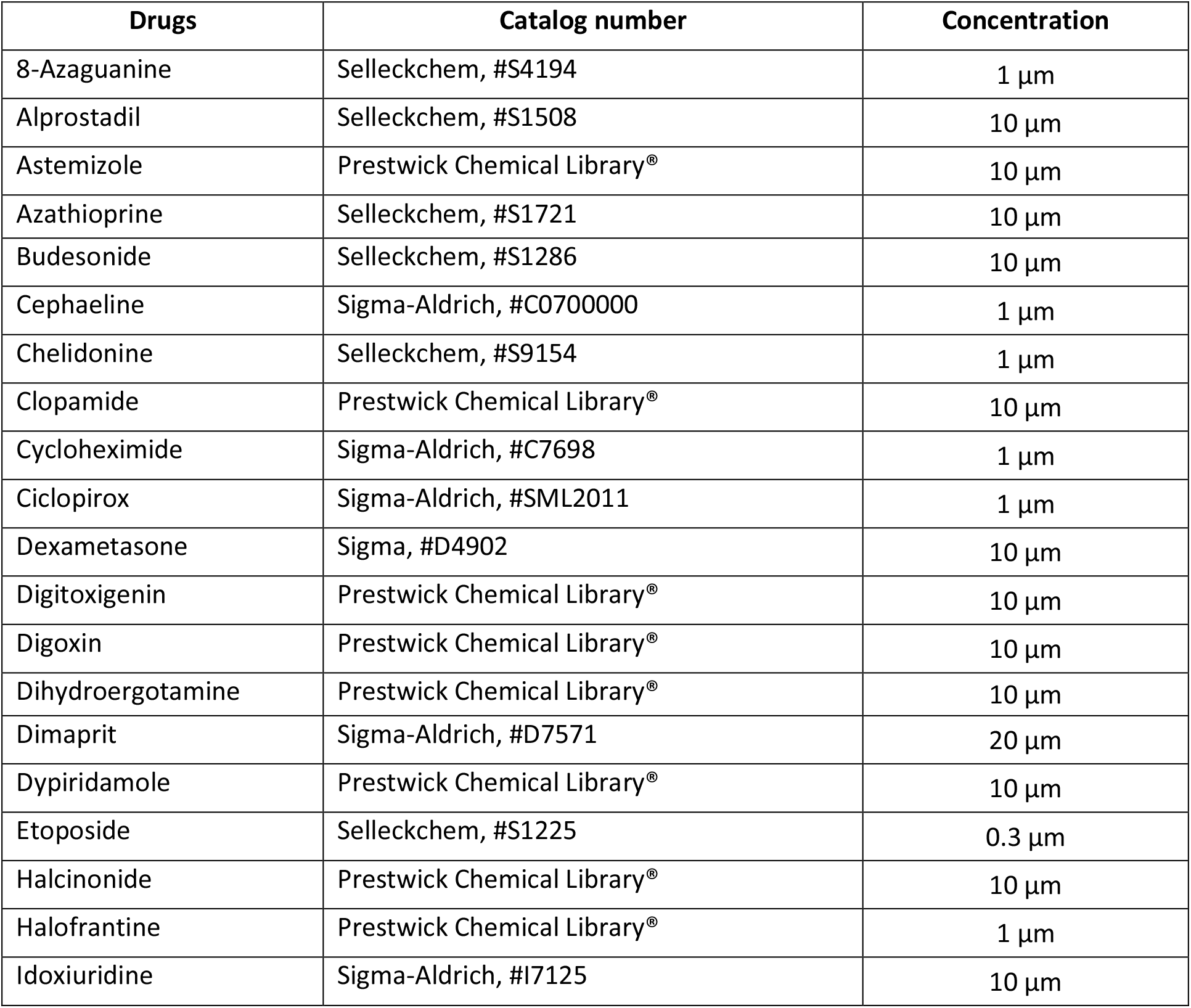
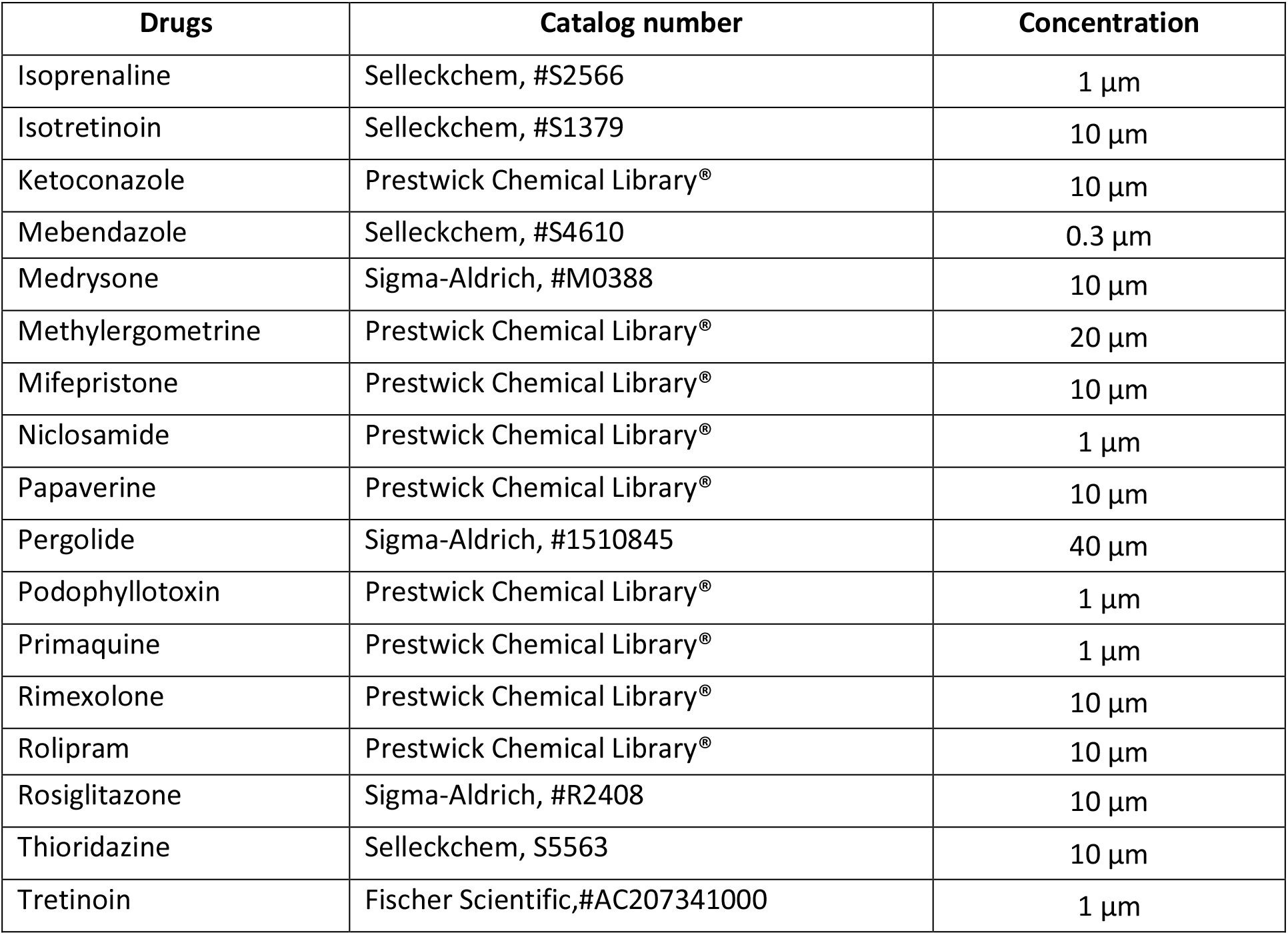
List of drugs used for experimental validation *in vitro*.

Drugs used for experimental validation were dissolved in dimethyl sulfoxide (DMSO) (Sigma-Aldrich, #D2650) or in alternative in Ethanol (Et-OH) in according to manufacturer’s instructions at a final concentration of 10 mM and stored at −20°C (see **Table 1**).

Compounds included in the commercial Prestwick Chemical Library^®^ (http://www.prestwickchemical.com), containing 1280 FDA-approved drugs, were pre-dissolved in 100% dimethyl sulfoxide (DMSO) at the final concentration of 10 mM and stored at −20 °C (see **Table 1**).

For monocytes/macrophage differentiation positive control, HL-60 were treated with PMA (Phorbol 12-myristate 13-acetate) (Selleckem, # S7791) 20 nM. DMSO 0.4% was used as negative control for all the experiments.

### 4.6 Cell viability

Cell viability was assessed by the trypan blue exclusion test. After 4 days of treatment, HL-60 cells were resuspended and mixed with Trypan blue solution dye (ThermoFisher, #T10282) 0.4%. When cell suspension is simply mixed with the dye could be visually examined to determine whether cells take up or exclude dye. Viable cells will have a clear cytoplasm whereas a nonviable cell will have a blue cytoplasm. Cell mixture was incubated for less than three minutes at room temperature and then was visualized using Countess™ II Automated Cell Counter (ThermoFisher, #AMQAX1000). For each sample percentage of viability cells has been collected and processed using GraphPad Prism 7 software.

### 4.7 Nitroblue Tetrazolium (NBT) assay

Nitroblue Tetrazolium (NBT) assay was performed after 4 days of drugs treatment. Following treatment with different compounds, cells were collected, centrifuged at 300 × g for 7 minutes at room temperature and resuspended in complete IMDM growth medium or in alternative in RPMI growth medium at the density of 3.0 × 10^5^ cells/mL with NBT (Nitro blue Tetrazolium Chloride) (Sigma-Aldrich, #N6876) 1mg/mL and MA (Phorbol 12-myristaPte 13-acetate) 5μg/ mL (Selleckem, # S7791). Next cells were incubated for 60 minutes at 37°C. In differentiated cells NBT is phagosomed, the intracellular enzymes convert NBT into insoluble blue formazan crystals. At the end of the incubation cells were collected, centrifuged at 300 × g for 7 minutes and washed in PBS (Dulbecco’s Phosphate Buffered Saline, Biowest, #L0625-500). Next cells were resuspended in complete IMDM growth medium or in alternative in RPMI growth medium and seeded on 12 well plate (Falcon^®^, #353043). For each sample images were acquired by light microscopy using an inverted microscope (Nikon, model Eclipse Ts2 #136710). At least 200 cells for sample were counted using ImageJ software. and the percentage of differentiated cells (cells containing blue-black formazan deposits) was calculated and processed using GraphPad Prism 7 software.

### 4.8 Flow cytometry

After 4 days of treatment HL-60 were resuspended, collected and centrifuged twice with PBS supplemented with BSA 0.5% (Bovine Serum Albumin, AppliChem, #A1391) and EDTA 2 mM at 300 × g for 10 minutes at 4°C. Cells pellet were then resuspended in PBS 2 mM EDTA 0.5% BSA and stained with antibodies (see **Table 2**) at the concentration of 1 × 10^6^ cells/ml for 30 minutes at 4° C. After incubation cells were washed in 1mL of PBS 0.5% BSA 2 mM EDTA and centrifuged 300 × g for 10 minutes at 4°C. Pellets were resuspended in 1 mL of PBS 2 mM EDTA 0.5% BSA. Stained cells were visualized on Cytoflex S, 3 lasers (488 nm, 405 nm and 638 nm) and 13 detectors. (Beckman Coulter). Live cells were gated based on side scatter and forward scatter. Approximately 10,000 events per samples were acquired. Quality control of the cytometer was assessed daily using CytoFLEX Daily QC Fluorospheres (Beckman Coulter, #B53230). Data were collected by CytExpert (Beckman Coulter) software. If needed, a compensation matrix was calculated using VersaComp Antibody Capture Kit (Beckman Coulter, #B22804) according to manufacturer’s instructions. FCS files were analysed using CytExpert software and the percentage of positive cells. In order to increase the throughput of samples, we used the plate loader of Cytoflex S for the acquisition. In this event cells were resuspended, collected and seeded on 96 well plate (Falcon^®^, #353072). Next, cells were stained with antibodies in 96 well plate as described above and finally 10,000 events per samples were acquired using a plate loader in Cytoflex S.

**Table 2:** Flow cytometry antibodies used.

| Flow cytometry antibodies | Concentration |
| --- | --- |
| PB450 anti-human CD11b (Bear1, #B16891) Beckman Coulter | 2 µg/mL |
| FITC anti-human CD71 (YDJ1.2.2, #IM0483) Beckman Coulter | 2 µg/mL |
| APC anti-human CD66b (80H3, #B15091) Beckman Coulter | 2 µg/mL |
| PC5.5 anti-human CD33 (D3HL60.251, #B36289) Beckman Coulter | 2 µg/mL |
| PB450 anti-human CD38 (LS198-4-3, #B92396) Beckman Coulter | 2 µg/mL |

### 4.9 Statistics

All the experiments have been conducted in at least 3 independent replicates derived from 3 cell line batches (n = 3). Data are presented as means ± standard error of the mean (SEM). Statistical significance was assessed using the Student’s t-test, One-Way ANOVA or Two-Way ANOVA according to the data set. Differences were considered statistically significant when p-value < 0.05. Plots and statistical analysis were produced using the GraphPad Prism 7 software.

## Author Contributions

Conceptualization, F.R., G.C.; methodology, F.R., E.M., G.C., C.F; formal analysis, F.R., E.M., R.S.; investigation, F.R., E.M., with the contribution of R.S., G.G., S.V., G.M., C.F.; writing—original draft preparation, F.R. E.M., G.C., C.F., G.G.; resources, G.C.; writing—review and editing, F.R., E.M., G.C.; supervision, G.C., L.C. C.F.; funding acquisition, G.C. All authors have read and agreed to the published version of the manuscript.

