## Supplementary Materials for "Transcription Factor Activation Profiles (TFAP) identify compounds promoting differentiation of Acute Myeloid Leukemia cell lines"

**Supplementary Table 1:** Annotation of Haematopoietic transcription factors

| TFs | Gene Name | Description | Reference (PMID) |
|---|---|---|---|
| CCAAT/enhancer-binding protein alpha | *CEBPA* | Promotes granulocytic differentiation | 11242107 |
| E3 SUMO-protein ligase EGR2 | *EGR2* | Promotes myeloid leukemia cell differentiation | 1864967 |
| Early growth response protein 1 | *EGR1* | Promotes myeloid leukemia cell differentiation | 1864967 |
| Endothelial transcription factor GATA-2 | *GATA2* | Required for myeloid differentiation | 12433372 |
| Runt-related transcription factor 1 | *RUNX1* | Required for hematopoiesis | 17431401 |
| Transcription factor AP-1 | *JUN* | Required for myeloid differentiation | 8423806 |
| Transcription factor PU.1 | *SPI1* | Controls myeloid and lymphoid differentiation | 23868921 |

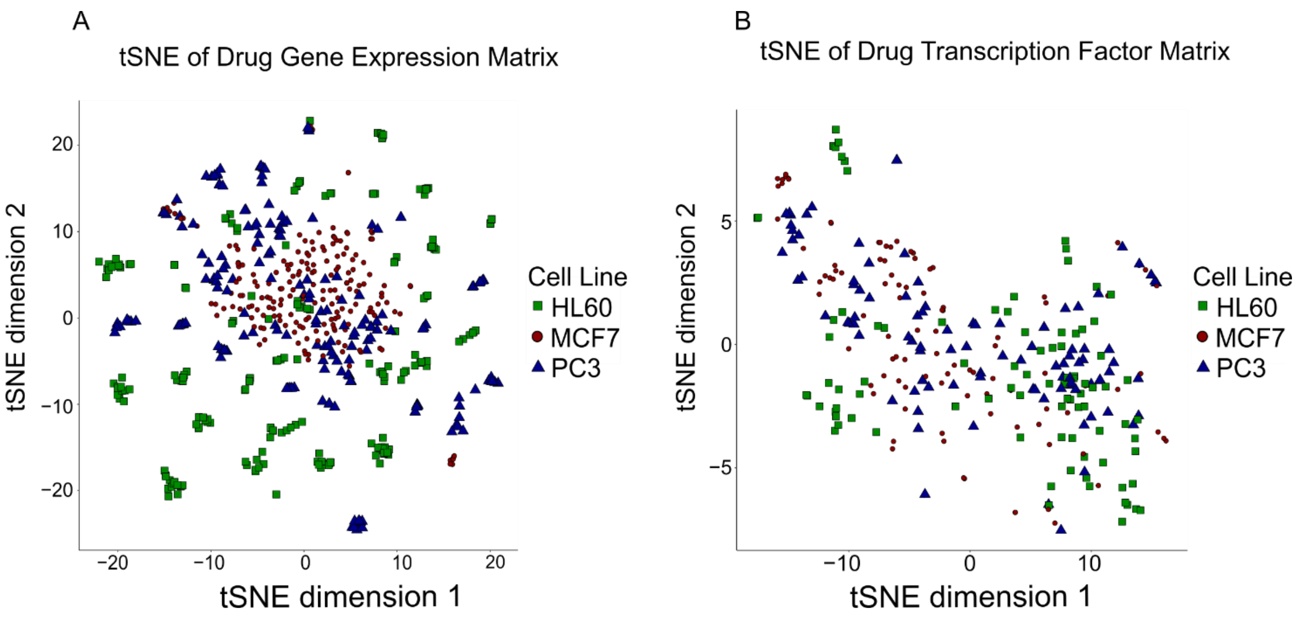


**Supplementary Figure 1**. tSNE maps of the multidimensional transcriptional profiles of different cell lines are incubated with different drugs. Each point correspond to different drug. Cell lines are represented with different shape and color. **(A)** tSNE map of gene expression profiles. **(B)** tSNE map of transcription factor profiles.

A

| Selected drugs | Therapeutic use |
|---|---|
| 8 Azaguanine | Potential antineoplastic activity |
| Alprostadil | Erectile dysfunction |
| Cephaeline | Gastrointestinal disorders |
| Chelidonine | Anticancer and antiviral activity |
| Cycloheximide | Antifungal activity |
| Cyclopirox | Moderate onychomycosis |
| Digitoxigenin | Esophageal cancer |
| Digoxin | Atrial fibrillation |
| Dihydroergotamine | Migraine headache |
| Dimaprit | Histamine agonist |
| Etoposide | Chemotherapeutic agent |
| Isoprenaline | Heart block |
| Isotretionoin | Severe recalcitrant nodular acne |
| Mebendazole | Broad spectrum anthelminthic |
| Medrysone | Allergic conjunctivitis |
| Methylergometrine | Control of excessive bleeding |
| Niclosamide | Tapeworm infestation |
| Pergolide | Parkinson's disease |
| Podophyllotoxin | External genital warts |
| Primaquine | Malaria |
| Thioridazine | Schizophrenia and anxiety disorder |
| Tretinoin | Acute promyelocitic leukemia |

B

| Selected drugs | Therapeutic use |
|---|---|
| Astemizole | Rhinitis and conjunctivitis |
| Azathioprine | Rheumatoid arthritis |
| Budesonide | Crohn's disease, ulcerative colitis |
| Clopamide | Antihypertensive |
| Dexametasone | Bacterial infections |
| Dypiridamole | Postoperative thromboembolic |
| Halcinonide | Dermatoses |
| Halofrantine | Severe malaria |
| Mifepristone | Termination intrauterine pregnancy |
| Ketoconazole | Endogenous Cushing's syndrome |
| Papaverine | Impotence and vasospasms |
| Rimexolone | Ocular inflammation |
| Rolipram | Antidepressive |
| Rosiglitazone | Type 2 diabetes mellitus |

C

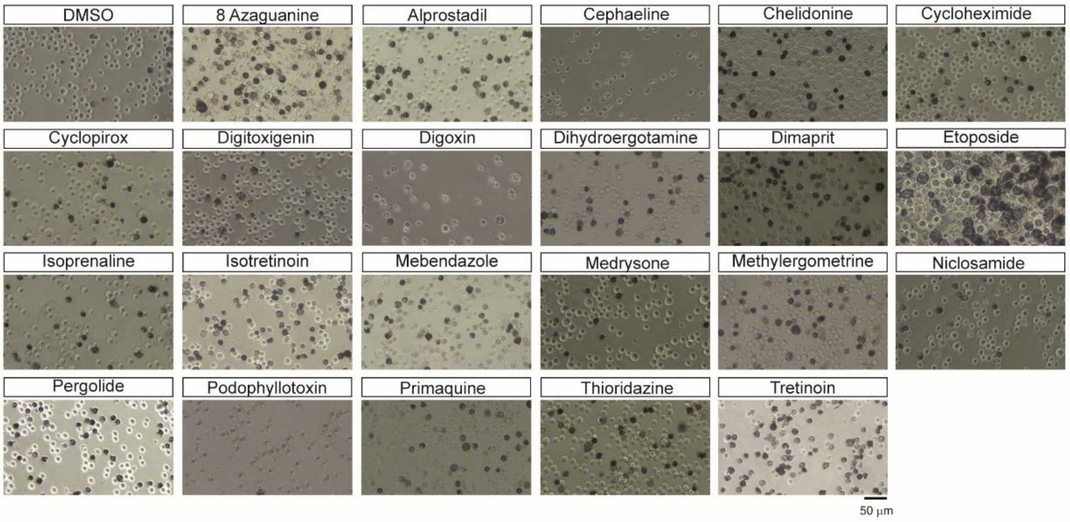


D

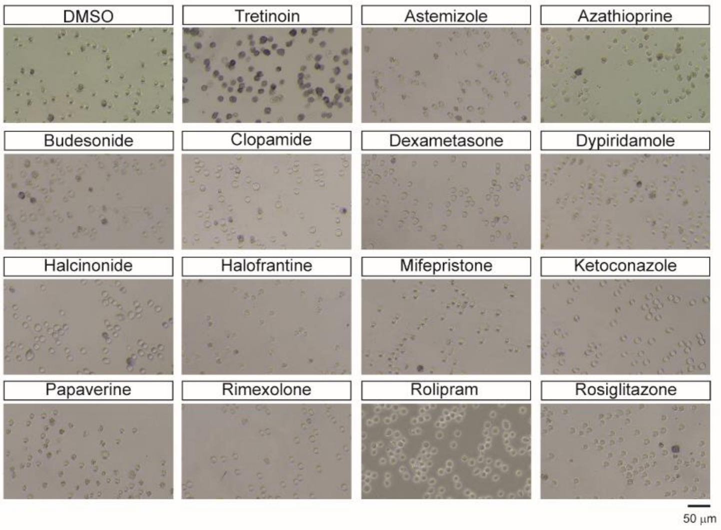


E

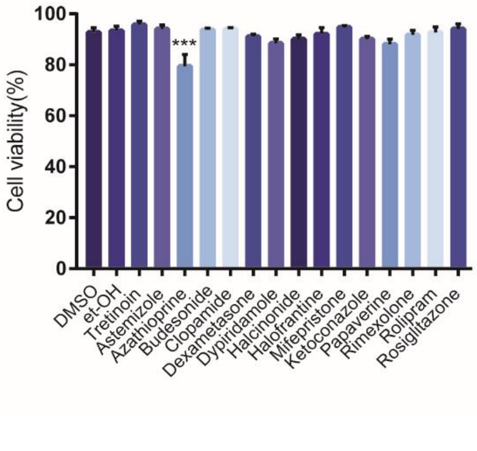

**Supplementary Figure2. (A)** List predicted drugs annotated with their therapeutic use. **(B)** List of the randomly chosen drugs that were tested for their ability to induce the differentiation of HL-60 cells. **(C)** Representative light microscope images of NBT staining in HL-60 after four days of the 22 selected drug treatment. Scale bar is 50 µm. **(D)** Representative light microscope images of NBT staining in HL-60 after four days of the randomly chosen drugs treatment. Scale bar is 50 µm. **(E)** Cell viability after four days of the randomly chosen drugs treatment. Cell viability was assessed by the Trypan blue exclusion test. Data are represented as means of three biological replicates ± SEM.Statistical analysis was performed using One-way ANOVA. Data are presented as mean ± SEM of three biological replicates. Significance * $p \leq 0.05$, ** $p \leq 0.01$, *** $p \leq 0.001$, **** $p \leq 0.0001$ and are related.
